# Comparative analysis of noise-attenuation mechanisms in gene expression: From single cells to cell populations

**DOI:** 10.1101/2023.04.06.535909

**Authors:** Zhanhao Zhang, Cesar Nieto, Abhyudai Singh

## Abstract

Negative feedback regulation is a well-known motif for suppressing deleterious fluctuations in gene product levels. We systematically compare two scenarios where negative feedback is either implemented in the protein production rate (regulated synthesis) or in the protein degradation rate (regulated degradation). Our results show that while in lownoise regimes both schemes are identical, they begin to show remarkable differences in high-noise regimes. Analytically solving for the probability distributions of the protein levels reveals that regulated synthesis is a better strategy to suppress random fluctuations while also minimizing protein levels dipping below a threshold. In contrast, regulated degradation is preferred if the goal is to minimize protein levels going beyond a threshold. Finally, we compare and contrast these distributions not only in a single cell over time but also in an expanding cell population where these effects can be buffered or exacerbated due to the coupling between expression and cell growth.

## I. Introduction

Gene expression is the process by which cells produce proteins from information stored in DNA. Due to the small number of molecules involved in this process and their short lifetimes, random fluctuations or noise in protein levels are an inherent feature of gene expression. Single-cell studies have shown that stochasticity in gene product levels is physiologically relevant, and impacts biological function often detrimentally, and in many cases beneficially [1]–[6]. For instance, while random fluctuations can enhance population adaptation to uncertain environments [7], they can be disadvantageous when homeostasis around a setpoint protein level is required [8]. Consequently, cells may evolve different mechanisms to suppress gene expression noise when it is detrimental [9], [10].

The gene expression process can be simplified as the contribution of two processes: protein synthesis and protein degradation (or dilution). When these processes reach equilibrium, protein concentration fluctuates randomly around a set point. As in engineering and physical systems, cells are known to implement negative feedback mechanisms to control the intensity of these fluctuations [11]–[16]. Simple negative feedback motifs are implemented when increasing intracellular levels of given protein results in a decreased synthesis rate, or alternatively, an enhanced degradation rate (Fig. 1A). These different strategies can exhibit distinct properties in terms of response time, efficiency, and stochastic behavior.

**Fig. 1.**
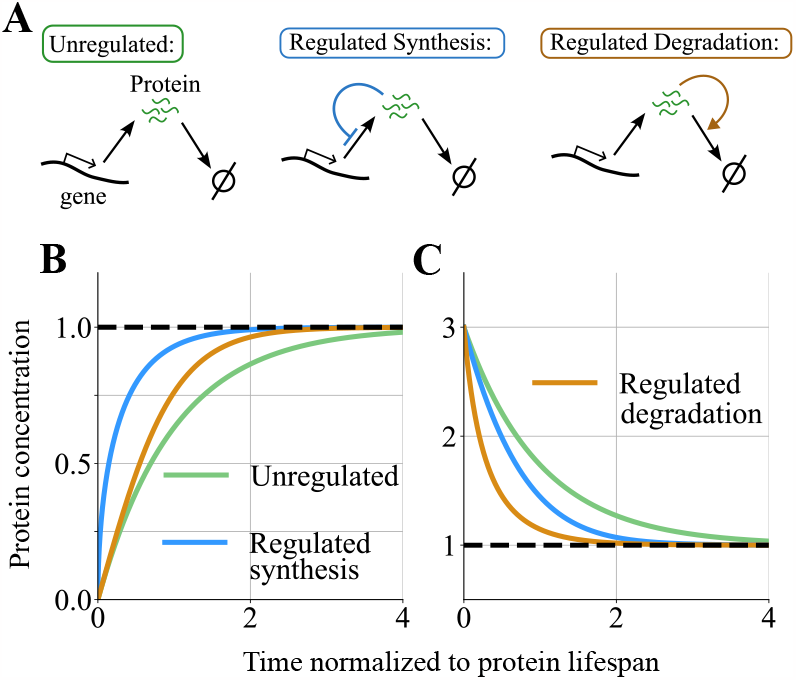
Negative feedback mechanisms in gene expression and their differential responses to large deviations from equilibrium. **(A)** The schematic of gene expression with and without feedback regulation. In the unregulated system, protein synthesis and degradation occur at given rates. When the synthesis is regulated, the higher the protein concentration the slower the proteins are produced. In self-regulated degradation, the higher the protein concentration, the faster these molecules are degraded. The solution of the nonlinear ordinary differential equations (3)-(4) are plotted over time by assuming *k* = *γ* = 1 and *g*(*x*) = *x* for a starting initial condition of *x*(0) = 0 (**B**) and *x*(0) = 3 (**C**). These parameter choices ensure that all models (1), (3) and (4) (with and without feedback) have the same steady-state protein concentration of 1 arbitrary units. Both negative feedback mechanisms have faster responses compared to the unregulated case [17]. Depending on the direction of the large perturbation, either synthesis-based or degradation-based feedback can provide faster reversion to equilibrium.

Among the possible applications of negative feedback implemented through self-regulation of protein synthesis, we can include fast kinetic response [17], suppressing random fluctuations in protein concentrations [18], stabilizing gene expression against mutations [19], and protein synthesis on demand [20]. On the other hand, negative feedback through self-regulation of the degradation rate has found applications in the sharp triggering and adaptation of response to stress [21], and in improving molecular sensing [16].

Both negative feedback mechanisms have been compared mainly at the level of small noise approximation [22]. At this level, both mechanisms show equivalence in terms of protein statistics. However, in a regime of higher random fluctuations in protein concentration, we expect appreciable differences in the dynamics of noise suppression [23]. As a result, we expect differences in the profiles of gene expression between the two feedback mechanisms with increasing intrinsic noise, such as making protein synthesis more burst-like rather than constitutive. To understand this divergence, we explore regimes beyond the small noise approximation using a combination of analytical approaches, and stochastic simulations of gene expression in both single cells and in proliferating cell populations.

We start by introducing a simple one-dimensional linear system to capture the dynamics of protein levels and transform it into a nonlinear system by having either the synthesis rate, or the degradation rate be protein-regulated. Next, we consider a stochastic formulation of this system, where protein synthesis occurs in random bursts of activity, and feedback is implemented in either the frequency with which bursts occur or the rate of protein degradation in between two consecutive bursts. The simplicity of this system results in an exact analytical solution for the steady-state distributions, which we contrast between the two feedback strategies in both small and high-noise regimes.

## II. Deterministic formulation of self-regulation mechanisms

To get a basic understanding of the gene expression process, let us simplify it as a deterministic system following the first-order differential equation

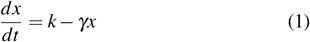

where the positively-valued scalar *x*(*t*) is the protein concentration at time *t* with steady-state

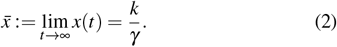

Here *k* is the synthesis rate that combines both transcription/translation processes and *γ* is the rate at which proteins are degraded. A key assumption underlying this onedimensional model is that the corresponding mRNAs are short-lived and are degraded rapidly compared to *γ* [24].

Let *g*(*x*) be a positively-valued continuously-differentiable function that monotonically increases with *x*. Then, negative feedback control can be independent in two alternative and orthogonal ways increasing protein concentration either decreases the synthesis rate

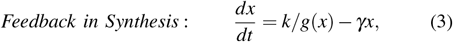

or enhances the degradation rate

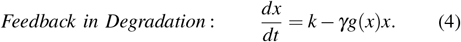

We further assume that

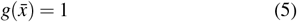

which ensures the exact same unique equilibrium point 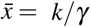 for both feedback scenarios.

It is straightforward to see that for small perturbations

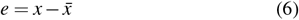

around the equilibrium point, linearizing nonlinearities in feedback models (3) and (4) yields the same linear system

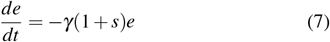

where

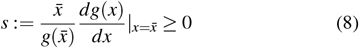

is the dimensionless log-sensitivity of *g*(*x*) with respect to *x* evaluated at the equilibrium point. While the dynamics of feedback-regulated synthesis and degradation are identical for small perturbations, nonlinearities result in different responses for large perturbations negative feedback in synthesis provides a faster response when starting from a low concentration (Fig. 1B). In contrast, degradation feedback is faster in reverting the system back to equilibrium when starting from a high initial concentration (Fig. 1C). These response kinetics to small and large perturbations are critical in determining the extent and shape of concentration fluctuations in the stochastic model, where noise constantly “kicks” the system out of equilibrium.

## III. Stochastic formulation of self-regulation mechanisms at low-noise regime

From this section onward, the concentration *x*(*t*) will be a positively-valued random process, the angular bracket notation ⟨ ⟩ will denote the expected value of random variables/processes, and we redefine 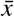 to be the steady-state mean concentration.

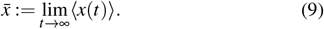

### A. Unregulated gene expression

Given its analytical tractability, we consider protein synthesis to occur in random bursts. The burst size *B*_*x*_ corresponds to the sudden increase in protein concentration due to protein synthesis in the short mRNA lifespan. This *B*_*x*_ is assumed to be an independent and identically distributed random variable drawn from an arbitrary positively-valued probability distribution and mean ⟨*B*_*x*_⟩. In the absence of regulation, the burst events arrive according to a Poisson process with a rate *k*_*x*_ (burst frequency), and each burst increases protein concentration by *B*_*x*_. It can be assumed that the number of proteins is relatively high. Hence, we can approximate the dynamics of the protein, such as between successive bursts, the concentration continuously decays according to the first-order kinetics [25], [26].

These processes can be gathered in a Stochastic Hybrid System (SHS) that is conveniently represented by

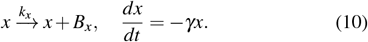

To quantify the extent of fluctuation in *x*(*t*) we use the tools of moment dynamics – for the SHS (10) the time evolution of uncentered moments is given by

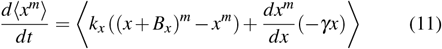

for any integer *m* ∈ {1, 2, …} [27], [28]. Simplifying (11) for *m* = 1 and *m* = 2 yields

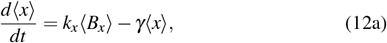

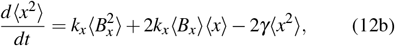

where for the equivalence of (12a) to the deterministic model (1) we need the net synthesis rate *k* = *k*_*x*_ ⟨*B*_*x*_⟩.

Solving this system of differential equations at steady-state

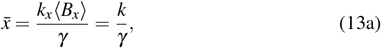

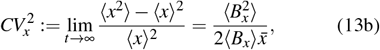

where the latter equation quantifies the magnitude of noise in protein concentration as determined by its steady-state coefficient of variation *CV*_*x*_.

### B. Self-regulated synthesis

Within this framework of bursty expression, negative feedback control of protein synthesis can occur both in the burst frequency or burst size [29]. Here, we consider feedback on the arrival of bursts, and alter the burst frequency to *k*_*x*_*/g*(*x*). Recall that as *g*(*x*) is a monotonically-increasing function, the arrival of bursts slows down with increasing protein concentrations. The moment dynamics for this case is similar to (11) with *k*_*x*_ now replaced by *k*_*x*_*/g*(*x*),

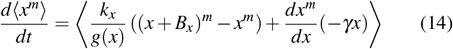

and these moment equations cannot be solved for arbitrary nonlinear functions *g*(*x*) due to issues with moment closure [30]–[35].

Towards that end, exploiting the Linear Noise Approximation (LNA) that considers small fluctuations of *x*(*t*) around the mean 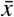 [36]–[38], one can linearize

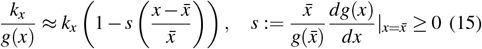

using the fact that 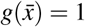. Now using this linear approximation in place of *k*_*x*_*/g*(*x*) in (14), and again performing the steady-state moment computations results in

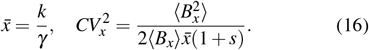

Note the noise levels here with feedback are lower by a factor of 1 + *s* as compared to the no-feedback case (13b).

### C. Self-regulated degradation

Negative feedback in degradation can be implemented by having a constant burst frequency and modifying the decay kinetics to

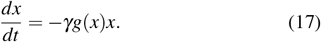

It is important to point out that if one considers a protein that is not actively degraded but its concentration is diluted from an increase in cell size (as is the case of most proteins in *E. coli*), this feedback implies a coupling between expression and cell growth. More specifically, lower expression of this protein will result in a lower exponential growth rate for cell size, resulting in longer cell-cycle times. We discuss some implications of this coupling in more detail in Section 5. With the altered nonlinear degradation, the moment dynamics is given by

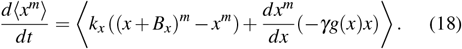

It turns out that linearizing *g*(*x*)*x* in (18) for small concentration fluctuations around the mean results in the exact same mean and noise levels as computed in (16) for feedback in synthesis. Thus consistent with previous observations [12], [39], *both feedback are identical in terms of noise attenuation for small fluctuations*.

### D. Inclusion of parametric fluctuations

Fluctuations in model parameters can be incorporated by, for example, replacing parameter *k*_*x*_ in the above modes by 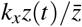, where *z*(*t*) is a stochastic process with mean 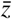 and coefficient of variation *CV*_*z*_. In the literature, such random variations in parameters are often referred to as extrinsic noise [40]–[44]. Analogous to the formulation of *x*(*t*) as an SHS in (10), *z*(*t*) can be modeling similarly with random bursts and exponential decay with rate *γ*_*z*_

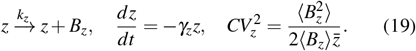

Repeating the analysis with the linear noise approximation for both types of feedback reveals the exact same noise levels

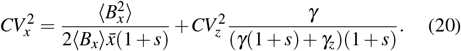

The second term on the right-hand side of (20) is the additional contribution from parametric fluctuations that attenuate much more precipitously with increasing *s* relative to the first term [45], [46]. In summary, the equivalence between the two feedback types in the small-noise operating regime extends to the inclusion of extrinsic noise.

## IV. Feedback comparison in high-noise regime

We now explore the stochastic feedback systems in the high-noise regime that correspond to large protein burst sizes. Throughout this section and later, we consider the physiological case of *B*_*x*_ drawn from an exponential distribution with mean ⟨*B*_*x*_⟩ [25], [47].

Our approach relies on analytically solving the steady-state distribution for the protein concentration and contrasting their shapes between feedback strategies as a function of ⟨*B*_*x*_⟩. As we increase ⟨*B*_*x*_ ⟩, we also correspondingly decrease the frequency with which bursts arrive to maintain the same protein concentration in the small-noise regime. As we will later see, given the nonlinearities of the system, the mean protein concentration in the stochastic model deviates considerably from its deterministic counterpart in the high-noise regime.

### A. Self-regulated synthesis

Recall that in this case, bursts arrive with a rate *k*_*x*_*/g*(*x*). Let *p*(*x, t*) denote the probability density function (pdf) of the protein concentration at time *t*. Thenm based on the SHS formalism introduced earlier, the time evolution of *p*(*x, t*) is described by the Chapman-Kolmogorov equation

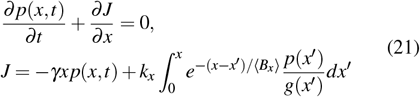

[25]. Assuming *k*_*x*_ = 1*/* ⟨*B*_*x*_⟩, *γ* = 1 and *g*(*x*) = *x* that ensures a steady-state mean protein concentration of 1 units in the small-noise regime for both regulated and unregulated systems, (21) has an exact analytical solution for the steadystate pdf

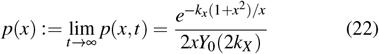

[39]. Here *Y*_0_(2*k*_*x*_) is the modified Bessel function of the second kind that can be computed in *Wolfram Mathematica* using

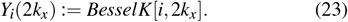

The steady-state pdf (22) leads to the following mean and the noise levels

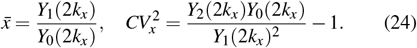

### B. Self-regulated degradation

The complementary case of feedback in degradation has exponentially-distributed bursts arrive with constant rate *k*_*x*_ = 1*/* ⟨*B*_*x*_⟩, and protein concentrations decay nonlinearly between consecutive bursts as

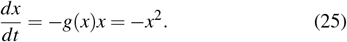

Formulating an analogous Chapman-Kolmogorov equation to (21) yields the following steady-state pdf

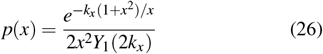

with the mean and the noise levels

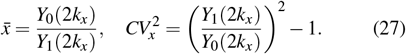

Fig. 2 presents a comparison of noise levels (24) and (27) with increasing mean bust size, together with unregulated noise levels (13b), and the LNA-predicted noise levels (16). For an exponentially-distributed burst size *B*_*x*_, *k*_*x*_ = 1*/* ⟨*B*_*x*_⟩, *γ* = 1 and *g*(*x*) = *x*, the unregulated and LNA-predicted noise levels are simply ⟨*B*_*x*_⟩ and ⟨*B*_*x*_⟩ */*2, respectively. It is quite evident in Fig. 2 that feedback regulation strategies have a considerable impact on noise suppression. As predicted earlier, noise levels for both feedbacks converge together with the LNA-predicted level in the low-noise regime. Conversely, in the high-noise regime, we see a divergence among them, with feedback in synthesis providing the lowest noise.

**Fig. 2.**
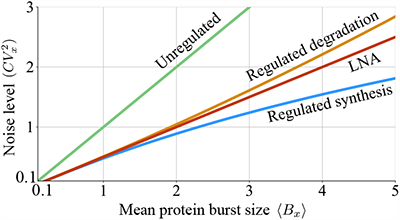
Self-regulated synthesis outperforms self-regulated degradation in attenuating noise at high protein burst sizes. The noise levels (24) and (27) obtained for self-regulated synthesis and degradation, respectively, are plotted with increasing mean protein burst size ⟨*B*_*x*_⟩. For comparison, noise level (13b) when there is no feedback, and LNA-predicted noise level (16) are also plotted. For this plot, *B*_*x*_ is drawn from an exponential distribution with mean ⟨*B*_*x*_⟩, *k*_*x*_ = 1*/*⟨*B*_*x*_⟩, *γ* = 1 and *g*(*x*) = *x*.

Fig. 3 shows sample trajectories of protein concentration evolving over time as well as the steady-state pdfs for different regulation strategies. In the low noise regime, characterized by ⟨*B*_*x*_⟩ = 0.1, the protein pdfs of both systems exhibit notable similarities. Meanwhile, Fig. 3B (bottom) presents the distributions of the two systems in the high-noise regime, where ⟨*B*_*x*_⟩ = 1. The distribution for the regulated degradation system demonstrates a faster decay of the tail (also evident in the 1*/x*^2^ scaling in (26) as compared to 1*/x* in (22)). The faster decay of the tail can be intuitively understood from the perturbation responses in Fig. 1 with quicker mean-reversion from a high concentration seen in degradation feedback as compared to synthesis feedback. Thus, in order to minimize concentration fluctuations above a high critical threshold, the regulated degradation approach is preferred. Using similar logic, it can be seen that in scenarios where the minimization of protein fluctuations below a low threshold is desired, the regulated synthesis strategy proves to be more effective.

**Fig. 3.**
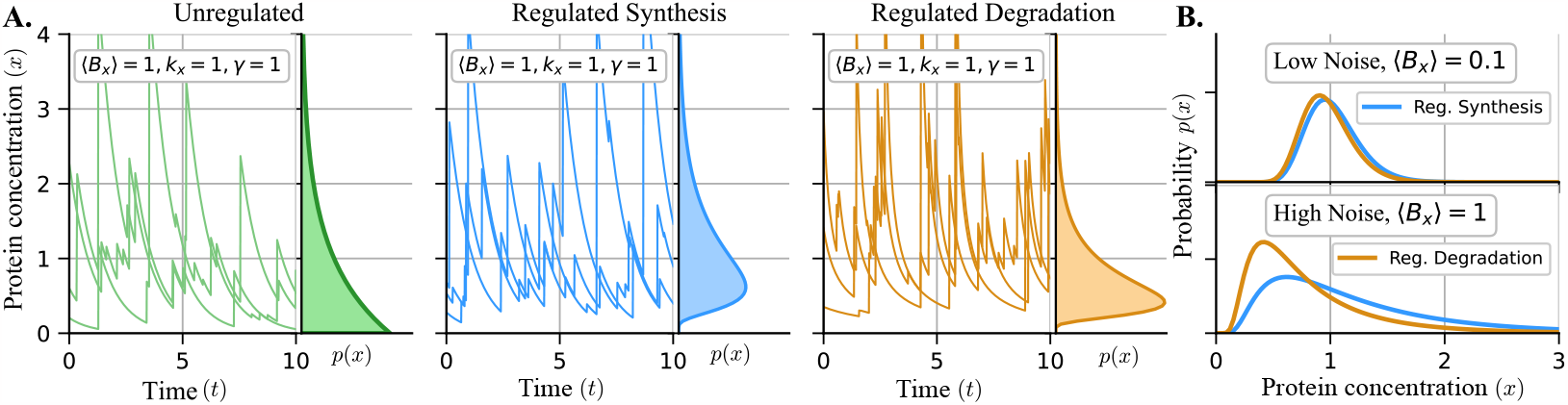
Self-regulated degradation leads to thinner-tailed steady-state protein concentration distributions compared to self-regulated synthesis. **A**: Sample trajectories for the protein concentrations for both unregulated and regulated gene expression with *g*(*x*) = *x* and *B*_*x*_ is drawn from an exponential distribution with the specified mean. **B**: The solutions of the steady-state protein distributions given by (22) and (26) demonstrate a close similarity in the low noise regime (top), where ⟨*B*_*x*_⟩ = 0.1, *k*_*x*_ = 10, and *γ* = 1. A noticeable deviation in the distribution of the two systems is observed in the high noise regime (bottom), where ⟨*B*_*x*_⟩ = *k*_*x*_ = *γ*_*x*_ = 1.

## V. Different perspectives for population statistics: Single-cell vs. population snapshots

Until now, this article has presented statistics of gene expression levels for independent cell trajectories. We will refer to this approach as the single-cell perspective. Another approach to the description of the process is to include the effects of cell proliferation [48], [49], as shown in Fig. 4. In the population-based approach, all descendants of a progenitor are considered, and the pdf is computed across all cells at a given time snapshot [50]–[52].

**Fig. 4.**
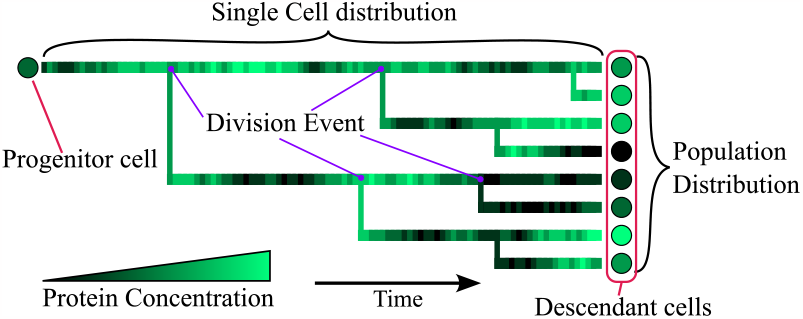
Comparing the statistics of protein concentration fluctuations between single-cell and population level. The schematic shows the expansion of the colony starting from an individual cell that proliferates to give birth to daughter cells. Statistics at the single-cell level can be interpreted as following a cell over time, and when it divides one of the two daughters is chosen randomly. At the population level, these statistics are estimated using all descendants of the progenitor cell at a given time point. Each circle represents one cell, and the color represents the protein concentration. A division occurs where the lineage of each cell is linked to another lineage.

We consider a long-lived protein whose concentration dilutes at a rate that is the same as the cellular growth rate. To obtain the population-level concentration distribution, we developed an agent-based model of a proliferating cell colony. If a cell dilutes its protein at a rate *γg*(*x*), then we consider division to be a stochastic process that occurs at the same rate, i.e., the probability that the cell divides during the next infinitesimal time interval (*t, t* + *dt*) is

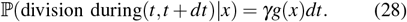

Just after a division event, a new cell is added to the population with the same protein concentration as the mother (neglecting any errors in the partitioning process). For self-regulated synthesis, the division rate of all cells is identical (*γ*). In this case, there is no difference in the analytically-obtained protein concentration pdf (22) versus the numerically obtained histogram from the population perspective (Fig. 5) both in lowand high-noise regimes.

**Fig. 5.**
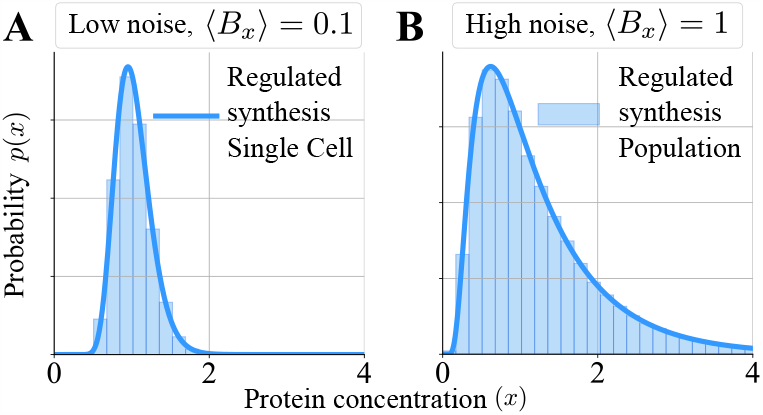
Single-cell and population-level distributions are identical for feedback in protein synthesis. The single-cell pdf from (22) and the histograms of the population distributions from simulation are plotted under a low noise regime (**A**), characterized by ⟨*B*_*x*_⟩ = 0.1, and a high noise regime (**B**), where ⟨*B*_*x*_⟩ = 1. *Parameters: γ* = 1, *k*_*x*_⟨*B*_*x*_⟩ = 1, and ⟨*B*_*x*_⟩ is marked on figures. *B*_*x*_ is drawn from an exponential distribution with mean ⟨*B*_*x*_⟩.

When the dilution is self-regulated, cells with higher protein concentrations are expected to replicate faster, and therefore contribute more towards the statistics. As illustrated in Fig. 6B, the population-level protein pdf in this case shows a noticeable right-shift compared to its single-cell counterpart. Fig. 6C plots the mean protein concentration against the burst size and all means converge to the same value of one arbitrary unit (as fixed earlier by parameter choice) for both feedbacks in single-cell and population perspectives. However, in the high-noise regime, the population-level mean for regulated dilution is higher than that of its single-cell perspective, and is closer to the mean for the regulated synthesis mechanism.

**Fig. 6.**
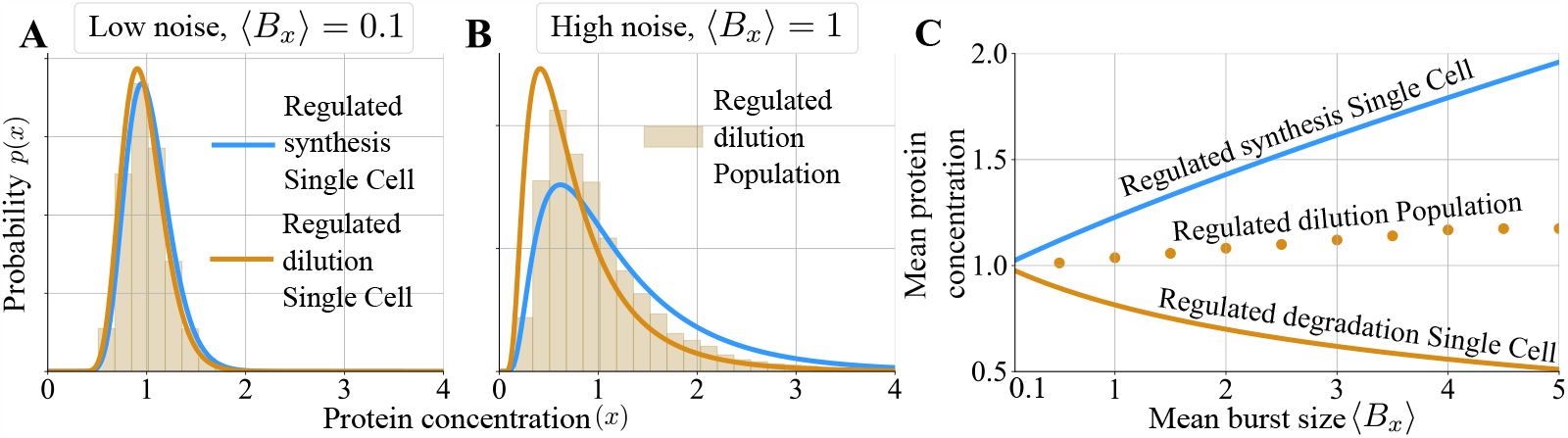
Negative feedback coupling between protein concentration and dilution drive differences in single-cell and population-level concentration pdfs. The single-cell pdf for the self-regulated dilution system (26) is compared to its population counterpart in the low noise regime (**A**), characterized by ⟨*B*_*x*_⟩ = 0.1, and a high noise (**B**), where ⟨*B*_*x*_⟩ = 1. For comparison purposes, the concentration pdf for self-regulated synthesis from Fig. 5 is also shown here. The mean steady-sate protein concentration calculated from the perspective of the single cell (24) and (27), and at the population level is shown as a function of the mean burst size ⟨*B*_*x*_⟩ (**C**). *Parameters: γ* = 1, *k*_*x*_⟨*B*_*x*_⟩ = 1, and the value of ⟨*B*_*x*_⟩ is depicted on each figure.

## VI. Discussion

In this contribution, we investigated stochastic dynamical systems motivated by the central question of noise attenuation in gene expression. Comparing two different feedback strategies of self-regulating protein synthesis versus degradation, we show that their perturbative linear dynamics around the set point are identical. This leads to the same noise level for both feedbacks when stochasticity is added either through bursty synthesis or parametric fluctuations.

As perturbations around the equilibrium begin to increase, these self-regulation mechanisms begin to diverge. At large protein burst sizes, regulated synthesis produces a lower *CV*_*x*_ as compared to feedback in degradation (Fig. 2). These differences are exemplified in the steady-state pdfs (Fig. 3) - *regulation in degradation results in a faster decay of the distribution tail*. Thus, regulated degradation would be a better strategy if protein levels that cross a high critical threshold are detrimental to cell fitness. In contrast, for an essential protein whose levels need to be maintained above a critical low threshold, negatively regulating protein synthesis is a better option. Similar observations have been made via stochastic simulation of genetic circuits [22], and here we quantify them analytically with exact solutions of the underlying Chapman–Kolmogorov equations.

Another key contribution of this work explores the steadystate protein level distribution across a cell population at a given time snapshot. In this population perspective, the protein distribution can differ from the distribution obtained by observing an individual cell over time. This is particularly relevant for stable proteins with no active degradation, where the protein dilution rate is essentially the cellular growth rate. *Our results show that both single-cell and population-level distributions are identical for self-regulated synthesis*, where cellular growth rates are constant (Fig. 5). However, when high protein levels increase the dilution and cell growth rate, cells with higher protein concentrations proliferate faster and become a larger part of the population. This results in a right shift of the population-level distribution as compared to its single-cell counterpart (Fig. 6). Based on these results, we expect a left-shifted population-level distribution for the case of positive feedback between expression level and cell growth rate. Positive feedback is implemented when high intracellular protein concentration slows the cellular growth rate, leading to lower dilution rate, which further increases the concentration [53]–[56].

It is interesting to note that the population-level distributions of both feedback strategies seem closer to each other, as compared to their single-cell distributions (Fig. 6). This leads to an interesting conclusion that negative feedback may suppress distribution differences at the population level, which in turn could be exacerbated by positive feedback where cellular growth is inversely related to expression levels. In summary, we have dissected the role of feedback in shaping the protein level distribution in single cells and in a proliferating population. Future work will consider mixtures of feedback in burst size, frequency, and degradation in combination with sequestration-based feedbacks [57]–[59] in determining the magnitude, time-scale, and exact shape of fluctuations in gene product levels.

